# Analysis of behavioral variables across domains and skin phenotypes in zebrafish: Role of brain monoamines

**DOI:** 10.1101/055657

**Authors:** Monica Gomes Lima, Anderson Manoel Herculano, Caio Maximino

## Abstract

Important neurochemical variations between strains or linages which correlate with behavioral differences have been identified in different species. Here, we report neurochemical and behavioral differences in four common zebrafish wild-type phenotypes (blue shortfin, longfin stripped, leopard and albino). Leopad zebrafish have been shown to display increased scototaxis in relation to the other strains, while both albino and leopard zebrafish show increased geotaxis. Moreover, leopard displayed increased nocifensive behavior, while albino zebrafish showed increased neophobia in the novel object task. Longfin zebrafish showed decreased turn frequency in both the novel tank and light/dark tests, and habituated faster in the novel tank, as well as displaying increased 5-HT levels. Leopard zebrafish showed decreased brain 5-HT levels and increased 5-HT turnover than other strains, and albino had increased brain DA levels. Finally, specific behavioral endpoints co-varied in terms of the behavioral and neurochemical differences between strains, identifying cross-test domains which included response to novelty, exploration-avoidance, general arousal, and activity.

## 1. Introduction

Important genetic variations between strains or lineages which correlate with behavioral differences have been identified in different species. The degree with which these variations explain the behavioral differences is not fully understood, but the use of behaviorally distinct strains might represent an important model to study the pharmacological and neurobiological correlates of behavior (Finn, Rutledge-Gorman, & Crabbe, 2003; Kalueff, Wheaton, & Murphy, 2007; Singewald, 2007; van der Staay, Arndt, & Nordquist, 2009). In this direction, mouse and rat inbred strains have been shown to differ in anxiety-like behavior and impulsivity (Brüske, Vendruscolo, & Ramos, 2007; Kangerski, Basso, Assreuy, Vendruscolo, & Takahashi, 2002; Neophytou et al., 2000; Ramos, Berton, Mormède, & Chaouloff, 1997; Trullas & Skolnick, 1993). The use of genetically tractable organisms, including invertebrate models and non-mammalian vertebrates, could generate important information regarding the genetic architecture underlying these disorders (Gerlai, 2010). For example, zebrafish from the leopard phenotype show lower serotonin levels in the brain and higher anxiety, a neurophenotype that is rescued by fluoxetine (Maximino et al., 2013), suggesting its use for pharmacological studies.

In this sense, the zebrafish (*Danio rerio* Hamilton 1822) represents important additions to this arsenal, presenting technical advancements such as optogenetics and transgenesis which facilitate the study of neural circuits (Friedrich, Jacobson, & Zhu, 2010; Rinkwitz, Mourrain, & Becker, 2011), complex behavioral phenotypes (Gerlai, 2010), and the presence of outbred strains which show considerable genetic variability (Coe et al., 2009). Inbred strains have been shown to differ in many important traits which are relevant for their behavior, such as brain transcriptome (Drew et al., 2012) and neurochemistry (Pan, Chaterjee, & Gerlai, 2012). Some of these strains have also been subjected to behavioral testing, such as the novel tank test, and shown to differ in terms of anxiety-like behavior (J. Cachat et al., 2011; Drew et al., 2012; Kiesel, Snekser, Ruhl, & McRobert, 2012; Maximino, Puty, Oliveira, & Herculano, 2013; Quadros et al., 2015; Sackerman et al., 2010; R. Y.Wong et al., 2012) and associated functions, such as habituation to novelty (Stewart et al., 2013; K. Wong et al., 2010) and risk-taking (Moretz, Martins, & Robison, 2007; Oswald & Robison, 2011). In zebrafish, inbred and outbred strains (such as AB, WIK, SH, Tü, and TL) exist which show some neurobehavioral differences, but commonly found mutant phenotypes (including skin mutant phenotypes such as leopard, albino and longfin, which in principle can arise in any strain) were also shown to possess important neurobehavioral differences (Egan et al., 2009; Maximino, Puty, Oliveira, et al., 2013; Quadros et al., 2015). For example, Egan et al. (2009) demonstrated that, in relation to wild-type shortfin, albino and leopard zebrafish show increased bottom-dwelling; Kiesel et al. (2012) demonstrated a similar profile for longfin mutants.

In the present work, we analyze the behavioral and neurochemical differences between the common WT phenotypes shortfin, longfin, leopard and albino. The novel tank and light/dark test were chosen as tests for anxiety-like behavior (Kysil et al., 2017); the novel object test was chosen as an assay for reactivity to novelty (Blaser and Heyser, 2015); and the nocifensive behavior test was chosen to measure sensory reactivity. Neurochemical endpoints involved the monoaminergic system, which has been implicated in defensive behavior in zebrafish (Maximino et al., 2016) and mammals (Morilak & Frazer, 2004).

## 2. Methods

### 2.1 Animals and housing

40 animals from the blue shortfin phenotype (‘shortfin’), 40 from the longfin stripped phenotype(‘longfin’), 40 from the albino phenotype (‘albino’) and 40 from the leopard pheno type (‘leopard’) were used in the present study. Animals were group-housed in mixed-phenotype 40 L tanks, with a maximum density of 25 fish per tank. Both male and female animals were used in the behavioral and neurochemical analyses. Tanks were filled with deionized and reconstituted water at room temperature (28 °C) and a pH of 7.0-8.0. Lighting was provided by fluorescent lamps in a cycle of14-10 hours (LD), according to the standard of care zebrafish (Lawrence, 2007). All manipulations minimized their potential suffering, as per the recommendations of the Conselho Nacional de Controle de Experimentação Animal (CONCEA, Brazil)(Conselho Nacional de Controle de Experimentação Animal-CONCEA, 2016). After behavioral experiments, one animal/phenotype/test was euthanized by immersion in cold water (T <4 °C) for at least 1 min {Formatting Citation}, and cessation of opercular movements was taken as a sign of death. Animals which were not euthanized were donated to private individuals.

### 2.2 Novel tank test

The protocol for the novel tank diving test (NTT) used was modified from Cachat et al.(2010) . Briefly, animals were transferred to the test apparatus, which consisted of a 15 × 25 × 20 cm (width × length × height) rectangular tank lighted from above by two 25 W fluorescent lamps, producing an average of 120 lumens above the tank. As soon as the animals were transferred to the apparatus, a webcam was activated and behavioral recording begun. Animals (n = 10 from each phenotype) were tested individually. The webcam filmed the apparatus from the front, thus recording the animal’s lateral and vertical distribution. Animals were allowed to freely explore the novel tank for 6 minutes. Video files were later analyzed by experimenters blind to the treatment using X-Plo-Rat 2005 (http://scotty.ffclrp.usp.br), and the images were divided in a 3 × 3 grid composed of 10 cm^2^ squares. The following variables were recorded:

– *time on top:* the time spent in the top third of the tank;
– *geotaxis habituation:* calculated as single-minute habituation rates (SHR), defined as the modulus of the difference between the time on top in the sixth minute and in the first minute;
– *squares crossed:* the number of 10 cm^2^ squares crossed by the animal during the entire session;
– *turn frequency:* the number of turns, defined as a change in more than 45° in the direction of swimming;
– *freezing:* the total duration of freezing events, defined as complete cessation of movements with the exception of eye and operculae movements;
– *homebase:* For the establishment of homebases, the number of visits and time spent in each 10 cm^2^ quadrant were calculated and expressed as % of total; a zone qualified as a homebase based on the maximal percentages for individual animals.

### 2.3 Light/dark test

Determination of scototaxis (light/dark preference; LDT) was carried as described elsewhere (Araujo et al., 2012; Maximino, de Brito, Dias, Gouveia Jr., & Morato, 2010; https://wiki.zfin.org/pages/viewpage.action?pageId=98537687). Briefly, animals were transferred to the central compartment of a black and white tank (15 cm X 10 cm X 45 cm h X d X l) for a 3-min. acclimation period, after which the doors which delimit this compartment were removed and the animal was allowed to freely explore the apparatus for 15 min. Animals (*n* = 10 from each phenotype) were tested individually. The following variables were recorded:

– *time on the white compartment:* the time spent in the top third of the tank (percentage of the trial);
– *squares crossed:* the number of 10 cm^2^ squares crossed by the animal in the white compartment;
– *latency to white:* the amount of time the animal spends in the black compartment before its first entry in the white compartment (s);
– *entries in white compartment:* the number of entries the animal makes in the white compartment in the whole session;
– *turn frequency:* the number of turns, defined as a change in more than 45° in the direction of swimming;
– *freezing:* the proportional duration of freezing events (in % of time in the white compartment), defined as complete cessation of movements with the exception of eye and operculae movements.
– *thigmotaxis:* the proportional duration of thigmotaxis events (in % of time in the white compartment), defined as swimming in a distance of 2 cm or less from the white compartment’s walls.
– *risk assessment:* the number of “risk assessment” events, defined as a fast (<1 s) entry in the white compartment followed by re-entry in the black compartment, or as a partial entry in the white compartment (i.e., the pectoral fin does not cross the midline).

### 2.4 Novel object exploration test

The novel object task (NOET) was adapted from Sneddon et al. (2003). Animals were transferred to a 15 × 25 × 20 (width × length × height) tank and allowed to acclimate for 5 minutes. After that period, a novel object (made up of a combination of red, yellow, green, blue and black Lego^®^ Duplo bricks such that the object was no longer than 9 cm in length and 6 cm in height) was slowly lowered into the tank (so as not to startle the fish) and placed at the center of a (previously defined) 10 cm diameter circle at the middle of the tank. Animals (*n* = 10 from each phenotype) were tested individually. A webcam filmed the apparatus from above, and the time spent within that circle and the number of squares crossed were recorded for 10 minutes.

### 2.5 Nocifensive behavior

To assess behavioral responses to a chemical, inescapable nociceptive stimulus, animals were acclimated to the test tanks (10 cm length X 10 cm width X 20 cm height Plexiglas tanks containing water from the home tank) for 30 min and then individual baseline (pretreatment) locomotor responses (number of 3 × 3 squares crossed during the session) were monitored for 5 min. Each fish was then individually cold-anesthetized and injected in the anal fin with a 1% solution of acetic acid. Afterwards, animals were returned to the original test tanks to recover from anesthesia, after which behavioral recording took place. Animals (*n* = 10 from each phenotype) were tested individually. The frequency of tail-beating events, in which the animal vigorously moves its tail but do not propel itself in the water (Maximino, 2011), and the change in total locomotion in relation to the baseline (Correia, Cunha, Scholze, & Stevens, 2011), were recorded as variables pertaining nocifensive behavior.

### 2.6 HPLC analysis of monoamines

Serotonin (5-HT), 5-hydroxy-indole-acetic acid (5-HIAA), norepinephrine (NE), dopamine (DA), 3,4-dihydroxyphenylacetic acid (DOPAC), 3-methoxy-4-hydroyphenylglycol (MHPG), and 3,4-dihydroxybenzylamine (DHBA) (50 mg) were dissolved in 100 mL of eluting solution (HClO_4_ 70% [0.2 N], 10 mg EDTA, 9.5 mg sodium metabissulfite) and frozen at-20 °C, to later be used as standards. The HPLC system consisted of a delivery pump (LC20-AT, Shimadzu), a 20 μL sample injector (Rheodyne), a degasser (DGA-20A5), and an analytical column (Shimadzu Shim-Pack VP-ODS, 250 × 4.6 mm internal diameter). The integrating recorder was a Shimadzu CBM-20A (Shimadzu, Kyoto, Japan). An electrochemical detector (Model L-ECD-6A) with glassy carbon was used at a voltage setting of +0.72 V, with a sensitivity set at 2 nA full deflection. The mobile phase consisted of a solution of 70 mM phosphate buffer (pH 2.9), 0.2 mM EDTA, 34.6765 mM SDS, 10% HPLC-grade methanol and 20% sodium metabissulfite as a conservative. The column temperature was set at 17 °C, and the isocratic flow rate was 1.6 ml/min. After the end of a specific behavioral test, animals were sacrificed and whole brains were dissected on ice-cold (< 4 °C) magnesium-and calcium-free phosphate-buffered saline (MCF) after sacrifice and homogenized in eluting solution, filtered through a 0.22 μm syringe filter, spiked with 0.22 μl of 2.27 mM DHBA (internal standard) and then injected into the HPLC system. These samples were prepared with a single brain per sample. These samples were derived from one animal per phenotype per test (totalling, thus, four brains per phenotype).

### 2.7 Statistical analysis

For continuous, normally-shaped variables (time on top of the novel tank, freezing, homebase, and SHR in the NTT; time on white, thigmotaxis, and freezing in the LDT; time near novel object in the NOET; activity reduction in the nocifensive behavior test; all neurochemical variables), data were analyzed via one-way analyses of variance (ANOVAs), followed by Tukey’s HSD when p < 0.05. Categorical variables (number of squares crossed and turn frequency in the NTT; number of entries in the white compartment, number of risk assessment events, and turn frequency in the LDT; number of squares crossed in the NOET; and tail beat frequency in the nocifensive behavior test) were analyzed via one-way Kruskal-Wallis’ tests, followed by Dunn’s test when p< 0.05. Latencies were analyzed by Mantel-Cox survival curves {Formatting Citation}. All analyses were made using GraphPad Prism 5.0.

In an attempt to expose the underlying structure of the many endpoints which were assessed, hierarchical clustering was applied to the data. Raw data were first transformed into Maximum Predictive Values (MPV), following the approach of Linker et al. (2011). Briefly, taking the data from wild-type shortfin animals as reference, for each variable the MPV was calculated as the ratio of the mean difference between two groups and their pooled standard deviations as follows:

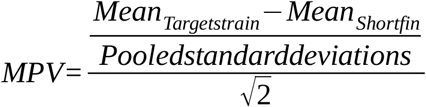

where

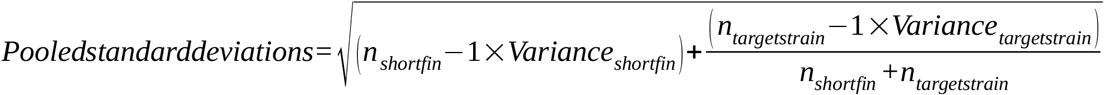

Given the mathematical simplicity of these measures, MPV scores were automatically calculated by LibreOffice Calc 3.6.6.2.

These scores represent the intensity (positive or negative) that fish from a given strain displayed a given behavioral endpoint in relation to shortfin fish. Resulting scores were normalized by centering each endpoint around the mean. Hierarchical clustering was then performed across behavioral endpoints and strains (‘arrays’) with Cluster 3.0 (University of Tokyo, Japan) using uncorrected correlation as clustering method, and single linkage as similarity metric. Clustering results were visualized as a dendrogram and colored “array” in Java TreeView (University of Glasgow, UK).

## 3 Results

### 3.1 Novel tank test

Albino and leopard fish spent less time in the top third of the novel tank than shortfin zebrafish (F_[3, 39]_ = 4.133, p = 0.0129; Figure 1A). No differences were observed between longfin zebrafish and other phenotypes. Longfin zebrafish crossed more squares in the 6-min. session than shortfin animals (p < 0.05), but no effects were observed between any other phenotype comparisons (H_[df = 4]_ = 7.622, p = 0.05; Figure 1B). In relation to longfin zebrafish, but not in relation to other phenotypes, leopard displayed greater turn frequency (H_[df = 4]_ = 17.45, p = 0.0006; Figure 1C). Longfin and albino froze more than shortfin, and leopard froze less than longfin (F_[3, 39]_ = 6.506, p = 0.0012; Figure 1D). Homebase behavior did not differ across phenotypes (F_[3, 39]_ = 0.9261, NS; Figure 1E). Habituation scores were higher in longfin than shortfin, smaller in albino than shortfin, and smaller in albino and leopard than in longfin (F_[3, 39]_ = 24.63, p < 0.0001; Figure 1F). With the exception of albino, fish from all phenotypes spent more time on the top in the last 3 min than in the first 3 min (F_[3,72]_ = 2.82, p = 0.0449; Figure 1G).

**Figure 1.**
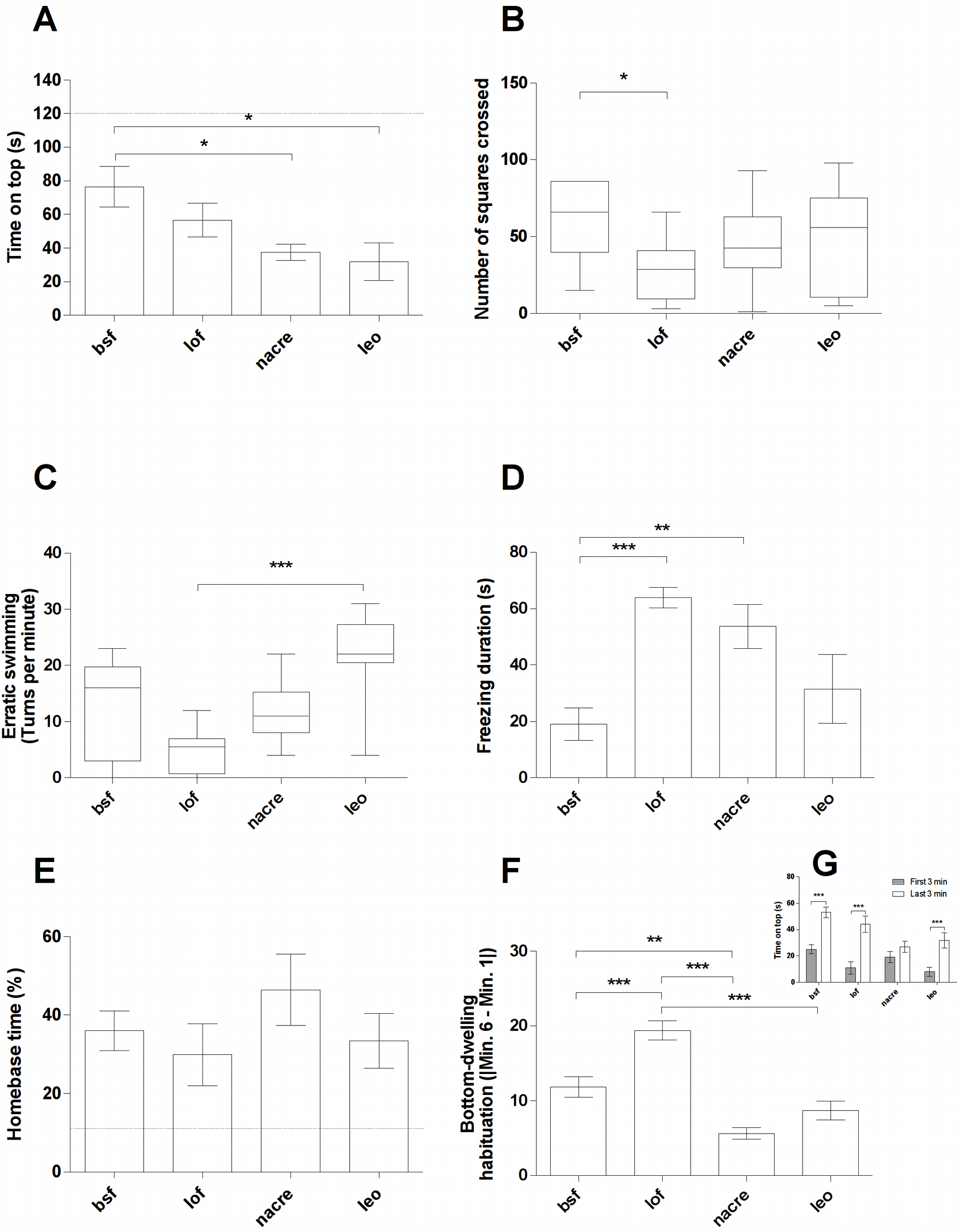
Behavioral differences between zebrafish from the blue shortfin (bsf), longfin (lof), albino(nacre), and leopard (leo) phenotypes in the novel tank test (NTT). (A) Time spent on the top third of the tank in the whole 6-min session; (B) Total number of squares crossed in the 6-min session; (C) Number of turns per minute; (D) Total freezing duration; (E) Time spent ina “homebase”; (F) Single-minute habituation score; (G) Time spent in the top third of the tank in the first 3-min (gray bars) and last 3-min (white bars). Bars represent mean ± standard error. Boxplots represent median ± interquartile range, with Tukey whiskers. ***, p < 0.001; **, p < 0.01; *, p < 0.05.

### 3.2 Light-dark test

Leopard zebrafish spend less time in the white compartment than shortfin and longfin zebrafish; no other phenotype differences were observed (F_[3, 39]_ = 12.66, p < 0.0001; Figure 2A). The latency to enter the white compartment (χ^2^ = 5.495, NS) or the number of entries in the compartment (H_[df = 4]_ = 1.414, NS) did not differ between strains (Figures 2B and 2C). Albino and leopard zebrafish showed increased risk assessment in relation to shortfin and longfin zebrafish (H_[df = 4]_ = 16.92, p = 0.0007; Figure 2D). No phenotype differences were observed in thigmotaxis in the white compartment (F_[3, 39]_ = 1.116, NS; Figure 2E). Leopard zebrafish displayed greater turn frequency in the white compartment than longfin zebrafish (H_[df = 4]_ = 9.561, p = 0.0227; Figure 2F). Finally, albino zebrafish froze more than shortfin, longfin and leopard zebrafish (F_[3, 39]_ = 16.20, p < 0.0001; Figure 2G).

**Figure 2.**
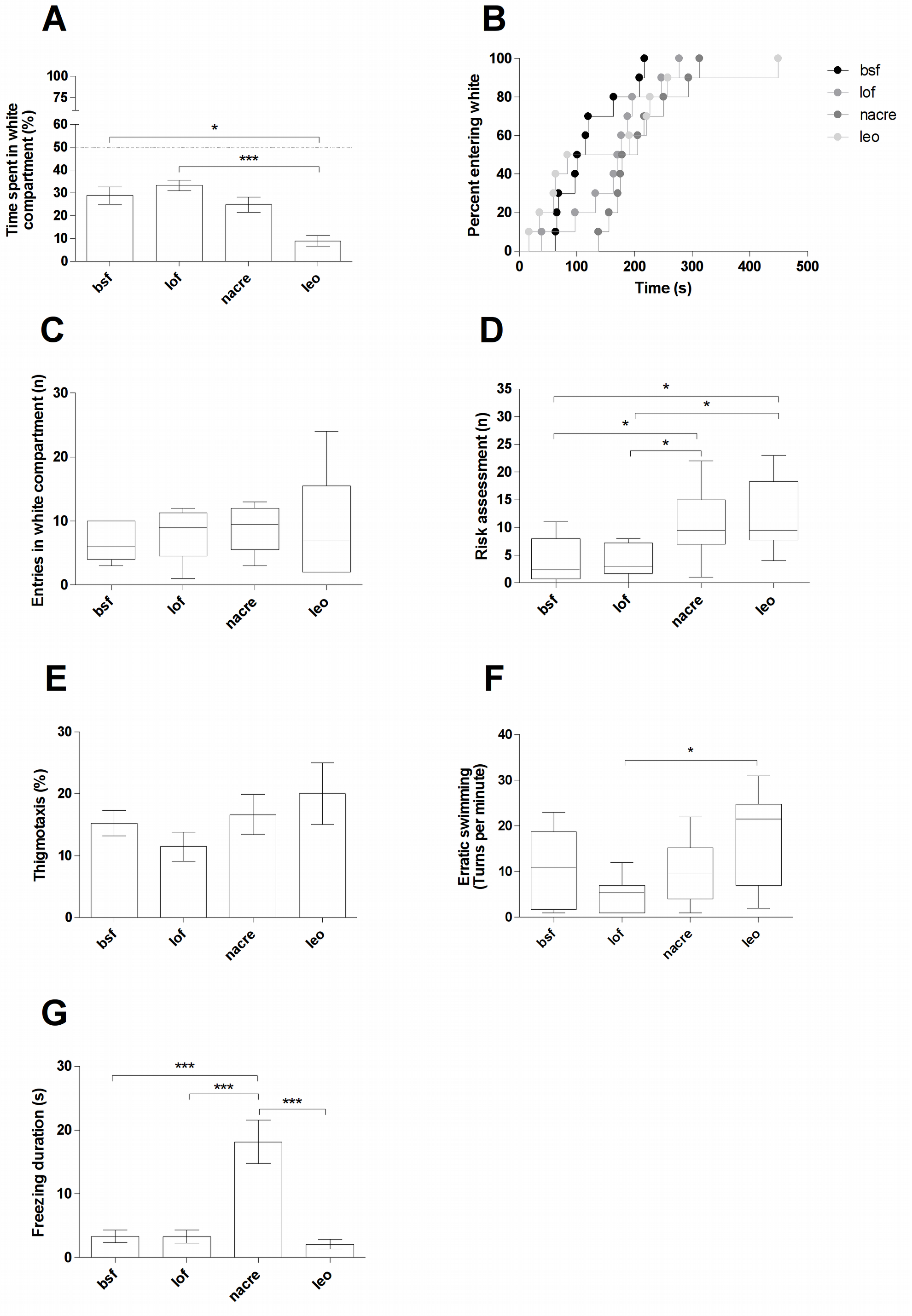
Behavioral differences between zebrafish from the blue shortfin (bsf), longfin (lof), albino(nacre), and leopard (leo) phenotypes in the light/dark test (LDT). (A) Time spent on the white compartment; (B) Latency to enter the white compartment; (C) Total number of entries in the white compartment; (D) Total number of risk assessment events; (E) Percent of the time on the white compartment spent in thigmotaxis; (F) Number of turns per minute on the white compartment; (G) Total duration of freezing in the white compartment. Bars represent mean ± standard error. Boxplots represent median ± interquartile range, with Tukey whiskers. Latencies are represented as Kaplan-Meier estimates of time until event. ***, p < 0.001; *, p < 0.05.

### 3.3 Novel object test

Albino zebrafish spent less time near the novel object than shortfin (F_[3, 39]_ = 3.918, p = 0.0161; Figure 3A). Locomotion in the novel object test was not different between phenotpypes (H = 6.406, NS; Figure 3B).

**Figure 3.**
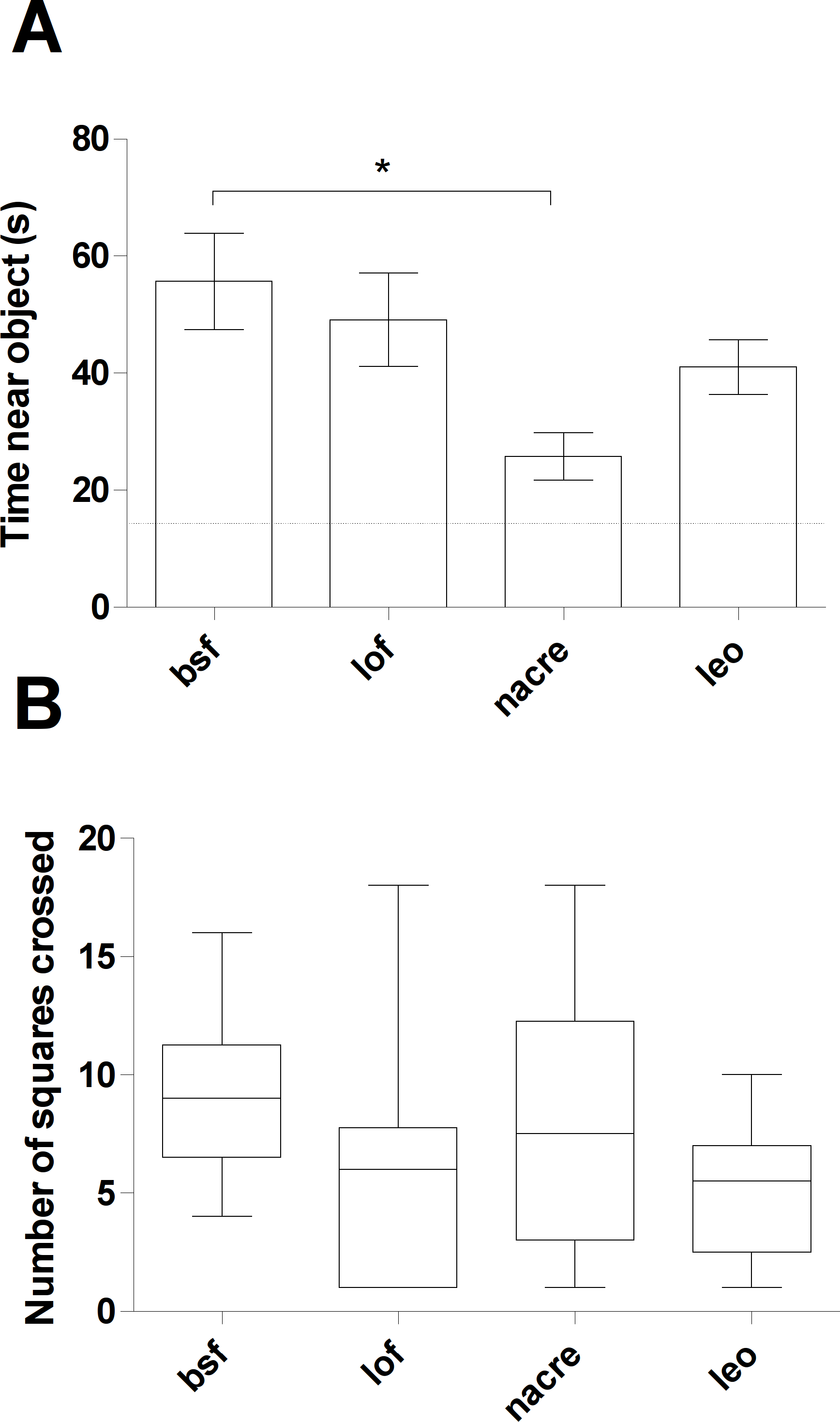
Behavioral differences between zebrafish from the blue shortfin (bsf), longfin (lof), albino(nacre), and leopard (leo) phenotypes in the novel object exploration test (NOET). (A) Time spent near the object in the whole 10-min session; (B) Total number of squares crossed in the 10-min session. Bars represent mean ± standard error. Boxplots represent median ± interquartile range, with Tukey whiskers. *, p < 0.05.

### 3.4 Nocifensive behavior

After injection of acetic acid in the tail, leopard zebrafish displayed more tail-beating than shortfin and albino animals (H_[df = 4]_ = 16.30, p = 0.001; Figure 4A). Likewise, leopard decreased their activity to a greater extent than shortfin zebrafish after this nociceptive manipulation (F_[3, 39]_ = 3.193, p = 0.035; Figure 4B).

**Figure 4.**
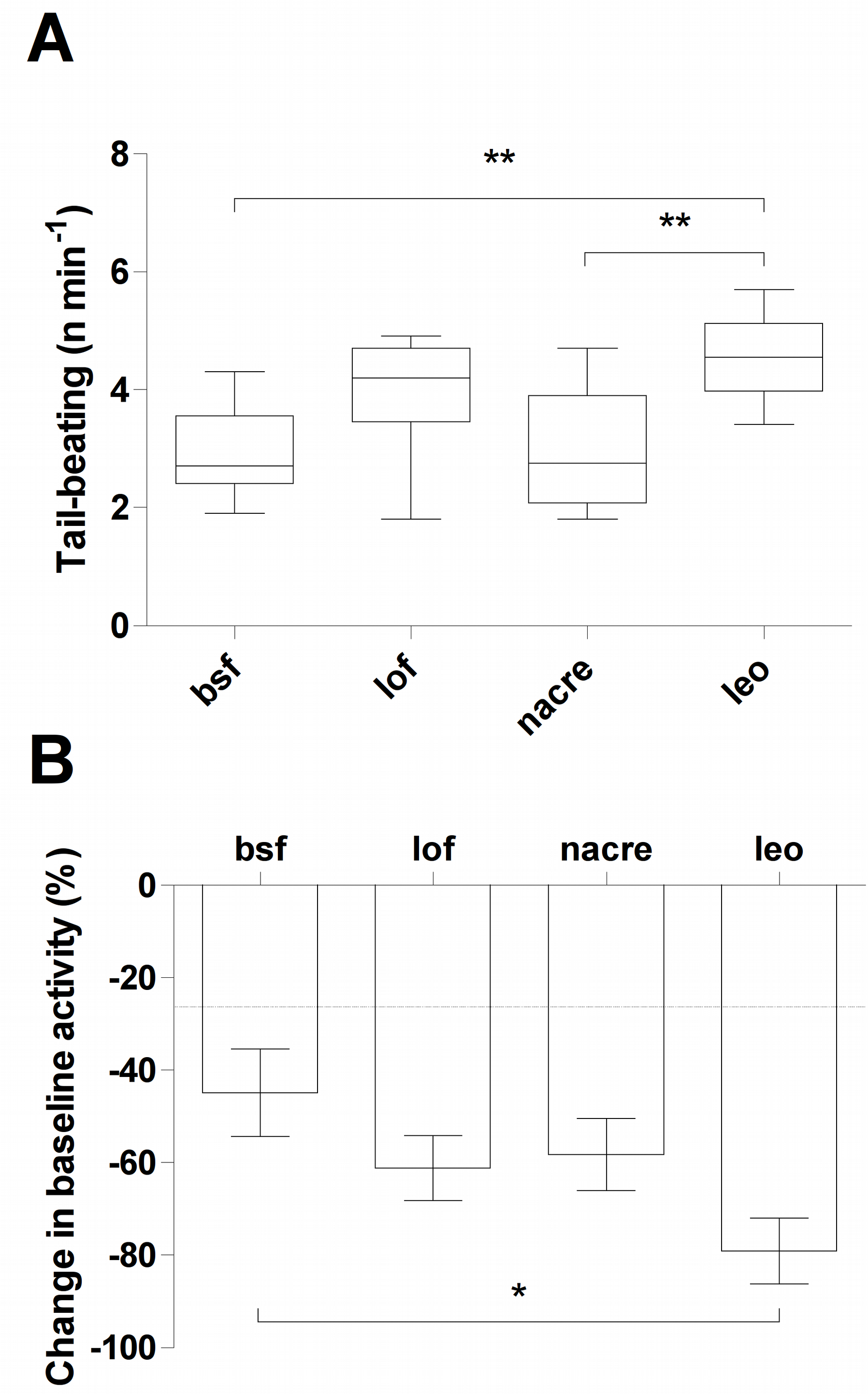
Behavioral differences between zebrafish from the blue shortfin (bsf), longfin (lof), albino(nacre), and leopard (leo) phenotypes in the nocifensive behavior assay. (A) Frequency of tail-beating events; (B) Change in baseline activity in relation to pre-injection levels. Barsrepresent mean ± standard error. Boxplots represent median ± interquartile range, with Tukey whiskers. **, p < 0.01; *, p <0.05.

### 3.5 Brain monoamines

Albino zebrafish showed higher brain dopamine levels in relation to shortfin animals (F_[3, 11]_ = 6.953, p = 0.0128; Figure 5A). No differences were observed in DOPAC levels (F_[3, 11]_ = 2.602, NS) or dopamine turnover rates (F_[3, 11]_ = 3.35, NS; Figures 5D and 5G). Longfin zebrafish had higher 5-HT levels than other strains, while leopard had lower levels than all phenotypes (F_[3, 11]_ = 21.94, p = 0.0003; Figure 5B). Longfin and leopard had lower 5-HIAA levels than other phenotypes (F_[3, 11]_ = 7.765, p = 0.0094; Figure 5E), and leopard had higher serotonin turnover than other phenotypes (F_[3, 11]_ = 12.16, p = 0.0024; Figure 5H). Shortfin animals had lower NE levels than longfin and leopard animals (F_[3, 11]_ = 8.198, p = 0.008; Figure 5C), while leopard had higher MHPG levels than all other phenotypes (F_[3, 11]_ = 12.06, p = 0.0024; Figure 5F). Finally, shortfin zebrafish had higher NE turnover rates than longfin animals (F_[3, 11]_ = 4.251, p = 0.0451).

**Figure 5.**
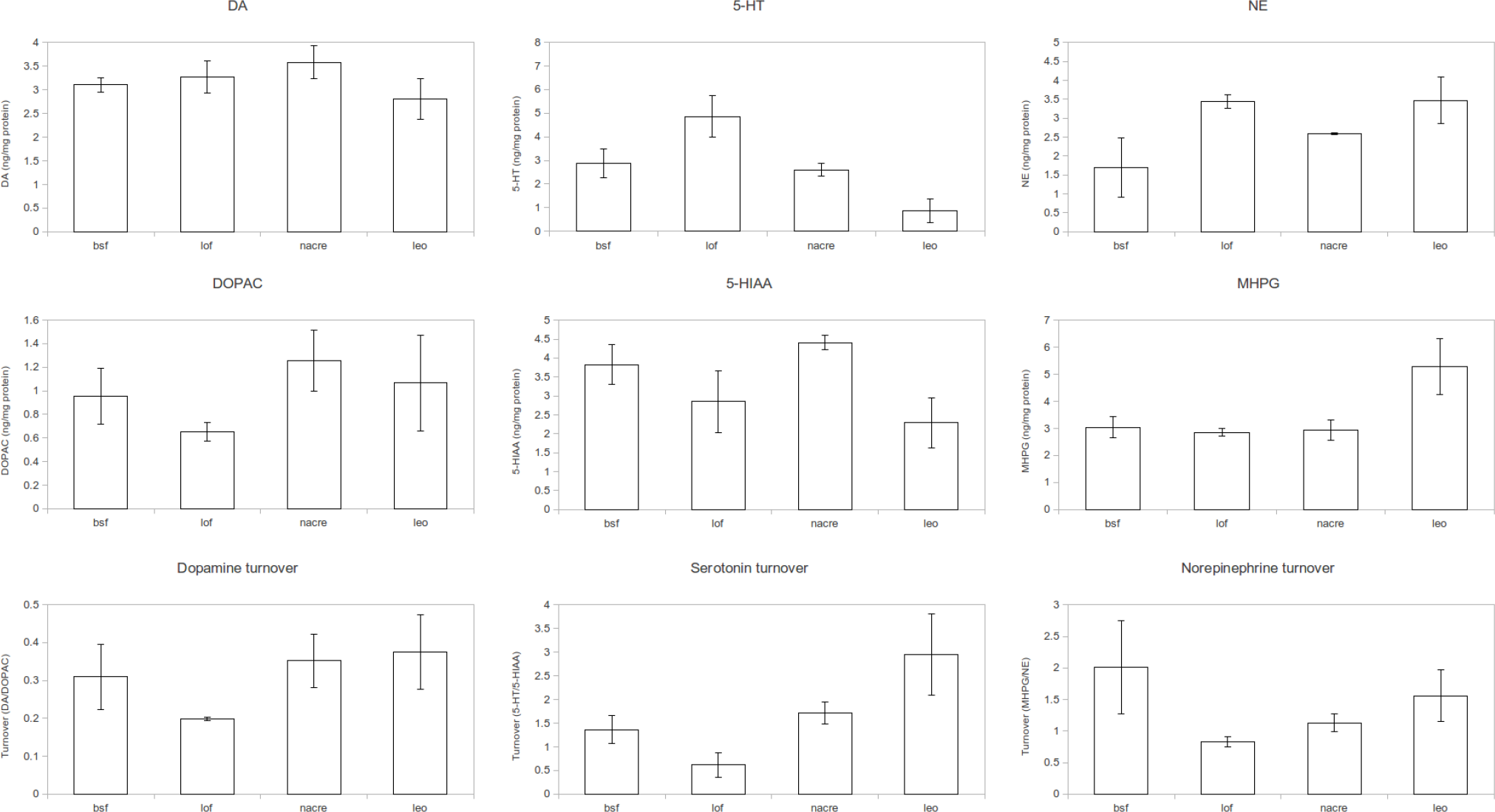
Monoamine levels in the brains of zebrafish from the blue shortfin (BSF), longfin (lof), albino (nacre), and leopard (leo) phenotypes. (A) Dopamine (DA) levels; (B) DOPAC levels; (C) Dopamine turnover (DOPAC:Dopamine ratios); (D) Serotonin (5-HT) levels; (E) 5-HIAA levels; (F) Serotonin turnover (5-HIAA:5-HT ratios); (G) Norepinephrine (NE) levels); (H) MHPG levels;(I) Norepinephrine turnover (MHPG:NE ratios). Bars represent mean ± standard error.***, p <0.001; **, p < 0.01; *, p < 0.05.

### 3.6 Clustering

Cluster analysis using longfin, albino and leopard zebrafish against a shortfin “reference” produced four identifiable behavior clusters (Figure 6). The first cluster (“response to novelty”) included tail-beating, time on top, habituation score, time near novel object and NE levels (r^2^ = 0.579); the second (“exploration-avoidance”) including freezing in the light/dark test, time on white, change in activity after acid injection, number of entries on white and DA and 5-HT levels (r^2^ = 0.741); the third (“general arousal”) including turn frequency in both the novel tank test and the light/dark test, locomotion in the novel tank test, thigmotaxis, risk assessment, latency to white, DOPAC and MHPG levels, and the turnover of all monoamines (r^2^ = 0.767); and the last (“activity”) including time spent on the homebase, locomotion in the novel object test, freezing in the novel tank test, and 5-HIAA levels (r^2^ = 0.930). Albino and leopard clustered together (r^2^ = −0.243), with longfin as outgroup. Raw MPV values can be found in Table 1.

**Table 1.** Calculated Maximum Predictive Values (MPVs) for the clustering analysis. Values in **bold** indicate statistically significant differences in post-hoc tests against shortfin animals(Figures 1–5).

| Test | Endpoint | Shortfin vs. |  |  |
| --- | --- | --- | --- | --- |
|  |  | Longfin | Albino | Leopard |
| NTT | Time on top | -0,295 | **-0,701** | **-0,633** |
|  | Turn frequency | -0,617 | -0,111 | 0,549 |
|  | Freezing | **1,529** | **0,833** | 0,218 |
|  | Locomotion | **-0,765** | -0,350 | -0,200 |
|  | Homebase | -0,153 | 0,235 | -0,070 |
|  | SHR | **0,831** | **-0,776** | -0,345 |
| LDT | Time on white | 0,234 | -0,183 | **-0,710** |
|  | Latency to white | 0,393 | 0,798 | 0,729 |
|  | Locomotion | 0,182 | 0,284 | -0,147 |
|  | Risk assessment | 0,013 | **0,640** | **0,751** |
|  | Thigmotaxis | -0,270 | 0,085 | 0,206 |
|  | Turn frequency | -0,434 | -0,077 | 0,388 |
|  | Freezing | -0,003 | **0,937** | -0,174 |
| NOET | Time near object | -0,134 | **-0,766** | -0,363 |
|  | Locomotion | -0,593 | -0,134 | -0,539 |
| Nociception | Tail-beating | 0,217 | 0,015 | 0,353 |
|  | Change in activity | -0,325 | -0,256 | **-0,680** |
| Neurochemistry | DA | 0,070 | **0,192** | -0,122 |
|  | DOPAC | -0,123 | 0,120 | 0,045 |
|  | DA turnover | -0,045 | 0,017 | 0,027 |
|  | 5-HT | **0,723** | -0,110 | **-0,770** |
|  | 5-HIAA | **-0,363** | 0,230 | **-0,582** |
|  | 5-HT turnover | -0,300 | 0,140 | **0,596** |
|  | NE | **0,663** | 0,340 | **0,653** |
|  | MHPG | -0,073 | -0,042 | **0,815** |
|  | NE turnover | **-0,453** | -0,337 | -0,170 |

**Figure 6.**
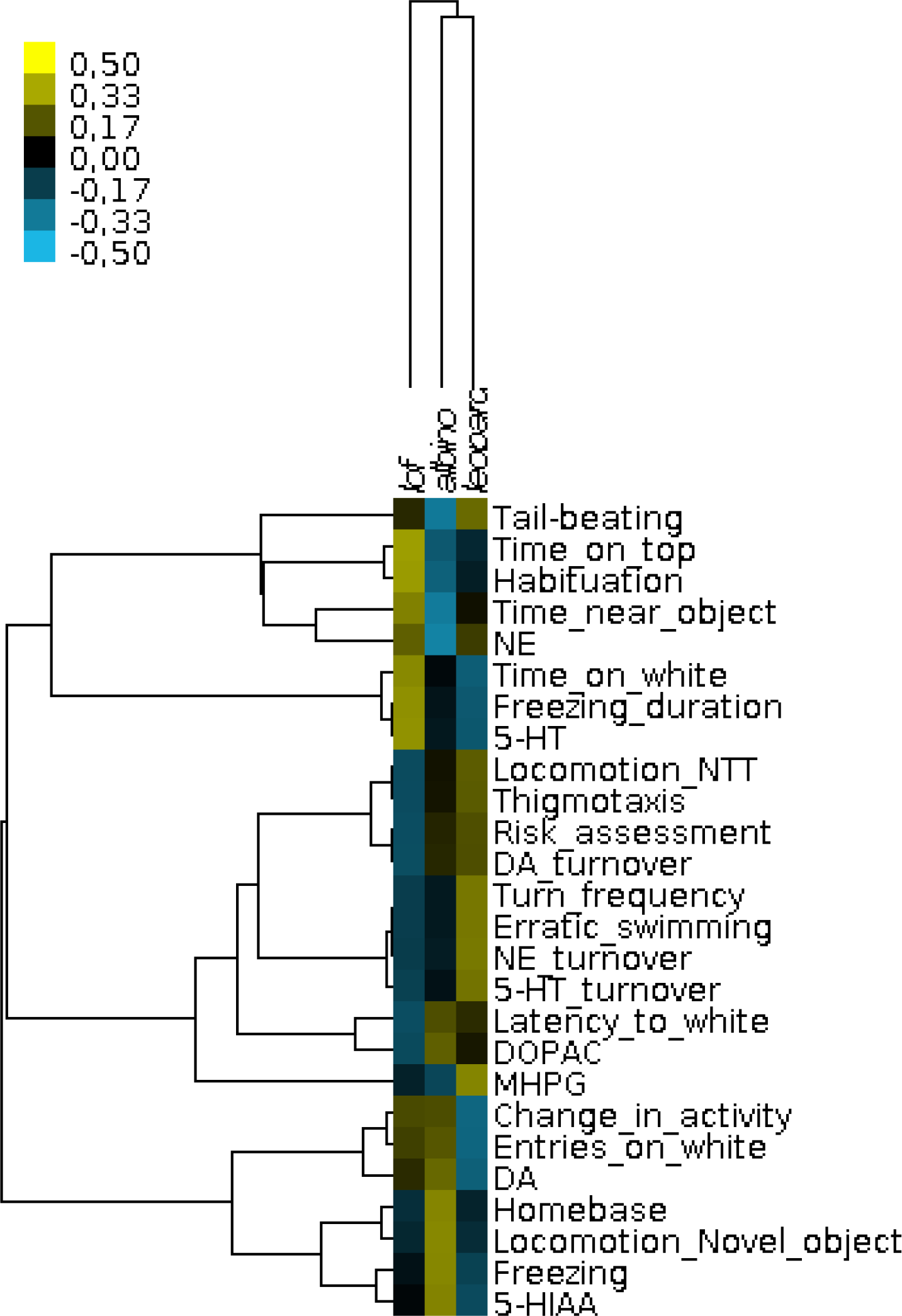
Hierarchical clustering of behavioral and neurochemical variables (rows) vs. phenotypes (columns). Clustering was made by calculating Maximum Predictive Values in relation to a reference phenotype (blue shortfin).

## 4. Discussion

In the present work, leopard zebrafish have been shown to display increased scototaxis in relation to the other phenotypes, while both albino and leopard zebrafish show increased geotaxis. Moreover, leopard displayed increased nocifensive behavior, while albino zebrafish showed increased neophobia in the novel object task. Longfin zebrafish showed decreased turn frequency in both the novel tank and light/dark tests, and habituated faster in the novel tank, as well as displaying increased 5-HT levels. Leopard zebrafish showed decreased brain 5-HT levels and increased 5-HT turnover than other phenotypes, and albino had increased brain DA levels. Finally, specific behavioral endpoints co-varied in terms of the behavioral and neurochemical differences between strains, identifying cross-test domains which included response to novelty, exploration-avoidance, general arousal, and activity.

In both the novel tank test and in the light/dark test, albino zebrafish showed increased freezing. While this behavior is poorly understood in zebrafish, freezing behavior does seem to vary with genetic background. Cachat et al. (2011) observed a small difference between shortfin and leopard zebrafish in freezing in the novel tank test, while no differences between longfin and leopard zebrafish were observed both in the NTT and the light/dark test (Maximino, Puty, Oliveira, et al., 2013). AB zebrafish selected for high freezing in the open field test show increased bottom-dwelling, increased alarm reaction, increased scototaxis and increased latency to feed in both disturbed and undisturbed conditions (R. Y. Wong et al., 2012). Blaser et al. (2010) demonstrated that zebrafish which consistently avoid the white compartment also freeze more after being confined to the white compartment. Thus, freezing seems to reflect either a fear response or a response to stressful manipulations, and therefore strains which show prominent freezing in the novel tank and light/dark tests could represent ‘reactive’ strains. Interestingly, the response of adult zebrafish with a mutation in the glucocorticoid receptor (*gr*^*s357*^) after transference to a novel environment is to freeze instead of explore, an effect which is reversed by acute diazepam or subchronic fluoxetine treatment (Ziv et al., 2012). Likewise, transient knockdown of *tyrosine hydroxylase 1* during development decreases freezing in adult zebrafish exposed to a novel tank (Formella et al., 2012), suggesting an important role for catecholamines in this response.

Lending credibility to such interpretations is the observation, in the present study, that albino zebrafish also show decreased exploration of novel objects, as well as decreased habituation in the novel tank test and increased risk assessment in the light/dark test. Differences between albino zebrafish and shortfin were observed in bottom-dwelling in the NTT (Egan et al., 2009). Thus, albinos appear to show enhanced stress responses to novelty, similarly to *gr*^*s357*^ mutants (Griffiths et al., 2012; Ziv et al., 2012). This response does not necessarily result from increased anxiety, as decreased ‘novelty-seeking’ could also be responsible for these results (Hughes, 1997, 2007). A non-selective increased responsiveness to stressors is discarded by the observation that albino display normal nociceptive behavior after acetic acid injection. Thus, this common mutant may represent an important addition in behavioral genetics in the sense that it shows slightly increased responsiveness to novelty, but not to nociceptive or simple anxiogenic stimuli.

In the literature, erratic or burst swimming has been defined as sharp changes in direction or velocity and repeated darting (Kalueff et al., 2013) which, in the novel tank test, are increased by ‘anxiogenic’ manipulations such as morphine withdrawal, alarm substance presentation and caffeine administration (J. Cachat et al., 2011); and decreased by acute fluoxetine and 5-HT_1B_ receptor antagonists (Maximino, Puty, Benzecry, et al., 2013). These measures commonly, but not necessarily, include the fast turns quantified in the present study as ‘turn frequency’. Turn frequencies are a way to quantify erratic swimming, an important behavioral endpoint sugggestive of anxiety-or fear-like states {Formatting Citation{; while this measure is not necessarily equivalent to erratic swimming measures reported elsewhere in the literature, it is of ecological relevance, because zebrafish can turn against the water current only until the current speed equals their routine maximum swimming speed (Plaut, 2000; Plaut & Gordon, 1994). In the present manuscript, turn frequency was higher in leopard than in other zebrafish phenotypes in both the novel tank and the light/dark test. Turn frequencies also did not differ between longfin and shortfin zebrafish in the present study, which should be expected if this variable was controlled solely by metabolic and/or biomechanic constraints, as observed in routine swimming (Plaut, 2000). Elsewhere, erratic swimming (and, presumably, turn frequency) was shown to not differ between shortfin and longfin zebrafish (Kiesel et al., 2012), or between shortfin and leopard (J. Cachat et al., 2011). Thus, the differences in erratic swimming between longfin and leopard found in the novel tank test and the light/dark test in the present work are likely to represent differences in anxiety and not metabolism or biomechanics.

From the neurochemical point of view, some observations call attention. First, all strains had higher norepinephrine levels than the reference shortfin. In the multivariate analysis, NE levels were grouped in the first cluster, which included tail-beating in the nocifensive behavior assay, time spent near the novel object, and time on top and habituation in the novel tank test. While NE has been proposed to mediate many different behaviors, in zebrafish noradrenergic drugs so far have been shown to modulate arousal (Ruuskanen, Peitsaro, Kaslin, Panula, & Scheinin, 2005). Along with increased responsiveness to sensory stimuli and voluntary motor activity, increased arousal leads to increased emotional reactivity (Pfaff, Martin, & Faber, 2012; Quinkert et al., 2011), and other neurotransmitter systems, including 5-HT (Cheng, Krishnan, & Jesuthasan, 2016; Yokogawa, Hannan, & Burgess, 2012), have been implicated in zebrafish arousal.

The third cluster extracted in our analysis shows behavioral endpoints more consistent with generalized arousal, such as turn frequency in both the novel tank test and the light/dark test, locomotion in the novel tank test, thigmotaxis, risk assessment and latency to white; moreover, the metabolites of dopamine and norepinephrine, DOPAC and MHPG, as well as the turnover rates of all neurotransmitters analyzed, clustered in this group.

While NErgic neurotransmission was higher in longfin, albino and leopard, 5-HT levels were lower in leopard, which also show increased anxiety-like behavior in the light/dark test and in the novel tank test, as well as increased nocifensive behavior. Leopard also showed increased 5-HT turnover and increased MPHG levels, suggesting increased monoamine oxidase activity. Dopamine levels were altered only in albino, reinforcing the hypothesis of elevated reactivity to novelty in these mutants. DA and 5-HT levels clustered together with change in activity in the nocifensive behavior assay, as well as freezing, time on white, and number of entries on white in the light/dark test. A role for serotonin in this assay has been proposed in zebrafish (Maximino, Puty, Benzecry, et al., 2013; Maximino, Puty, Oliveira, et al., 2013), and the 5-HT1A receptor was implicated in the antinociceptive effect of alarm substance (Maximino, Lima, Costa, Guedes, & Herculano, 2014), but so far little is known about the role of dopamine in scototaxis.

The heterogeneous nature of behavioral variation in this paper supports our anterior notion that behavioral tests of ‘anxiety’ in zebrafish do not necessarily measure the same dimensions (Maximino et al., 2012). The present results suggest that these tests fall under the aegis of ‘domain interplay’ (Kalueff, Ren-Patterson, LaPorte, & Murphy, 2008), with different behavioral endpoints mapping to different behavioral domains. Using a similar approach to cluster analysis presented in this paper, Cachat et al. demonstrated the existence of two major clusters in the novel tank test, the first including (among others) latency to upper half of the tank, freezing and erratic swimming, and the second including time spent in the upper half, distance traveled and average velocity. Importantly, these clusters grouped in relation to the effects of ‘anxiolytic’ manipulations (which decrease behaviors from the first cluster and increase behavior from the second) and ‘anxiogenic’ manipulations (with the opposite effect); the latter include animals from the leopard phenotype (J. Cachat et al., 2011). Moreover, clustering based on habituation rates, instead of anxiety level, produces different results in the same assay (Stewart et al., 2013), suggesting that anxiety and habituation are independent in the NTT. Leopard zebrafish also habituates freezing faster than shortfin, but this effect was inversely affected by exposure to an alarm substance or to acute caffeine treatment, and freezing habituation was actually increased by anxiolytic treatments (chronic ethanol, chronic fluoxetine, acute nicotine, chronic morphine) (Stewart et al., 2013).

In conclusion, the present paper demonstrated that common wild-type zebrafish phenotypes differ in their behavior in multiple behavioral assays, suggesting a genetic basis for conflict-and novelty-stress induced behavior, as well as in nocifensive behavior. Moreover, a monoaminergic substrate for these differences has also been described. In general, the identification of the genes and neural substrates underlying the behavioral variation of these common zebrafish mutants could represent important additions to the arsenal of tools to understand the neurogenetics of anxiety disorders.

## Acknowledgments

Part of this research was financed by a CNPq/Brazil grant (Process 483336/2009-2). AMH is recipient of a CAPES productivity grant.

